# Tumor suppressor p73 transcriptionally regulates *c-FLIP* to impede its priming of extrinsic apoptosis while a “switcher compound” degrades c-FLIP protein

**DOI:** 10.1101/2024.04.21.590479

**Authors:** Shengliang Zhang, Lanlan Zhou, Wafik S. El-Deiry

**Affiliations:** Laboratory of Translational Oncology and Experimental Cancer Therapeutics, Warren Alpert Medical School, Brown University, Providence, Rhode Island; Department of Pathology and Laboratory Medicine, Warren Alpert Medical School, Brown University, Providence, Rhode Island; The Joint Program in Cancer Biology, Brown University and Brown University Health System, Providence, Rhode Island; Legorreta Cancer Center at Brown University, Warren Alpert Medical School, Brown University, Providence, Rhode Island; Hematology/Oncology Division, Department of Medicine, Warren Alpert Medical School, Brown University, Providence, Rhode Island

**Keywords:** c-FLIP, p73, small molecule, mutant p53, cell fate, cancer therapy

## Abstract

The tumor suppressor p73 is a member of the p53 family and transcriptionally activates multiple p53-targets involved in cell cycle regulation and apoptosis. In addition to pro-apoptotic signaling, outcomes of p73 activation include cell survival signals. Thus, p73 activity and targets may provide insight in cell fate outcomes between cell survival and apoptosis following cellular stress. We report that *cellular FLICE inhibitory protein* (*c-FLIP*), a master antiapoptotic factor, is a transcriptional target of p73. The activation of p73 (α and β isoforms) transcriptionally upregulates *c-FLIP-L/S* expression in cancer cells. The cell fate decision following p73 activation is determined by the adjustment of the balance of outcomes of p73 activation between p73-induced pro-apoptotic signaling and *c-FLIP-L/S* expression in cancer cells. p73 primes extrinsic apoptosis via an autocrine death ligand-DR5 axis, and the priming appears to be titrated at the level of c-FLIP-L/S. The p73-upregulation of *c-FLIP-L/S* increases the threshold of extrinsic apoptosis. Cells with poor priming levels convert to cell cycle arrest and survival. Depletion of *c-FLIP-L/S* increases the p73-priming levels towards extrinsic apoptosis and sensitizes cancer cells to p73-primed extrinsic apoptosis. We further identified a small-molecule CB-7587351 (“switcher compound”) that alters p73 activation outcomes through c-FLIP-L/S protein degradation. Therapeutic activation of p73 can restore p53- signaling in mutant p53-expressing cancer cells effectively bypassing the p53 deficiency in cancer cells. Our discovery of p73 transcriptional upregulation of c-FLIP provides a promising strategy for depleting c-FLIP to improve antitumor efficacy of p73-targeting cancer therapy for p53-mutant tumors.

## Introduction

The tumor suppressor p73 is a family member of p53, a guardian of genome integrity through repair of DNA and regulation of the cell cycle. p73 and p53 have high structure and sequence homology. Like p53, p73 transcriptionally activates a subset of p53- targeted genes, leading to cell cycle arrest, cellular apoptosis and chemo-sensitivity [1, 2]. p73 loss contributes to tumor development [3–6]. Therefore, p73 is considered as a tumor suppressor. Unlike p53 which is frequently mutated in approximation of 50% tumors [7], p73 is rarely lost or mutated in cancer cells. Therefore, p73 appears to be a promising target to reinforce p53 pathway signaling for cancer therapy in p53-mutant or p53-null cancer cells [8]. Our laboratory previously reported that approximately half of the small molecules in the selected leads from the National Cancer Institute (NCI) diversity set library restored p53 pathway signaling through activation of p73 [9]. Small molecules, such as RETRA, Prodigiosin and NSC59984, activate p73 via releasing it from the inhibitory complex of mutant p53 [10–12]. These studies support a rationale for therapeutic activation of p73 to bypass and restore wild-type function to mutant p53 in cancer cells. However, there are no FDA approved p73-targeting compounds in clinical trials, and the therapeutic efficacy of p73 activators needs to be improved.

p73 has more than an anti-tumor effect as it has also been found to regulate development among other functions. p73 plays a critical role in maintaining neuron cell survival [13], and sustains cell stemness of cancer stem-like cells through redox metabolic reprograming [14]. A global gene expression analysis has shown that p73 activation included, but was not limited to, proapoptotic and survival signal patterns [15]. There is an unmet need for understanding how p73 determines the cell fate between cell survival and cellular apoptosis in response to cellular stresses. Answers to this question will be helpful to improve the antitumor efficacy of targeting p73.

Apoptosis is one of the major types of cell death and occurs through intrinsic and extrinsic pathways. The intrinsic pathway is initiated by intracellular stress through mitochondria, and the extrinsic pathway is triggered by external stimuli through cell death receptors [16]. The key factors involved in activating the two pathways, such as DR5 in the extrinsic pathway and PUMA in the intrinsic pathway, are transcriptional targets of p53 and p73 [8, 17]. Both pathways are considered as a predominant mechanism by which p53 and p73 induce cell apoptosis. Cellular FADD-like IL-1 beta-converting enzyme (FLCE)-inhibitory protein (c-FLIP) is an anti-apoptotic factor playing a key role in controlling the extrinsic apoptotic pathway by blocking the activation of caspase 8. The c-FLIP family comprises three isoforms at the protein level, c-FLIP-L (long isoform), c-FLIP-S (short isoform) and c-FLIP-R (Raji isoform), and all of them lack caspase activity therefore inhibit caspase 8 autoactivation when they are bound together [18, 19]. Recently, p53 and its other family member p63 both have been found to upregulate c-FLIP-L expression in cancer cells [20–22]. The p53-upregulation of c-FLIP-L partially leads to cancer cell resistance to Nutlin-mediated induction of p53- dependent apoptosis, but the resistance mechanism remains unclear [22]. A recent study reported a correlation between the increase of c-FLIP at the protein level and the overexpression of the TAp73α in hepatocellular carcinoma cells Hep3B and HepG2 [23]. There is a need to understand the role of the upregulation of c-FLIP in the cell fate decision within the p53 family to increase the antitumor efficacy of targeting the p53 family accordingly. In this study, we explored and describe the role of p73 (both α and β isoforms) as a transcription factor in upregulating c-FLIP-L/S expression in cancer cells. The p73-dependent upregulation of c-FLIP, in turn, results in poor priming of extrinsic apoptosis in cancer cells when p73 is activated. p73 appears pivotal in a cell fate decision by balancing its outcomes between c-FLIP-L/S expression and pro-apoptotic signaling in cancer cells. This insight was further exploited to identify a small molecule CB-7587351 as a “switcher compound” to adjust the outcome of p73 signaling through degradation of c-FLIP and activation of the p73 pathway in cancer cells. Our study develops a new strategy for harnessing c-FLIP-L/S depletion to increase the antitumor efficacy of targeting p73 in TP53-deficient cancer cells for cancer therapy.

## Materials and Methods

### High-Throughput Screening

Functional cell-based screening for small molecules that increase p53-transcriptional activity was performed as described in the previous study using noninvasive bioluminescence imaging in human colorectal cancer cells which stably express a p53 reporter, PG13-luc [10]. Briefly, cells were treated with 10 μM of compounds from a Chembridge library (50K small molecules), and p53 transcriptional activity was evaluated by bioluminescence from a p53-reporter in cells using an IVIS imaging system (Xenogen, Alameda, CA) at 2h, 24h and 72h after compound treatment. DMSO treatment was used as a negative control in each screened plate. Compounds that increased p53-responsive bioluminescence were selected for secondary screening.

### CellTiter-Glo luminescent Cell viability assay

Cells were seeded on 96-well plates with a density of 3000 cells/well and treated with small molecule compounds as desired for 72 hours. Cell viability was measured by CellTiter-Glo bioluminescence (Promega, catalog no. G7572) and analyzed using an IVIS imager.

#### Annexin V Apoptosis Assay

Cells were seeded on 96-well black plate and treated with the reagents as indicated in the figures. Annexin V levels were measured using RealTime-Glo™ Annexin V assay kit (Promega). The bioluminescent Annexin V was analyzed using an IVIS imager.

### Chromatin Immunoprecipitation (ChIP) PCR assay

ChIP was performed according to the protocol from Upstate Biotechnology with slight modifications as previously described [24]. Briefly, 2 × 10^6^ cells were fixed with 1% formaldehyde, followed by sonication which sheared the cross-linked DNA fragments to 200 – 1000 base pairs in length. The sonicated chromatin was incubated with 3 μg of anti-p73 antibody (A300-126A, Bethyl Laboratories) at 4°C overnight, and 50 μl of packed salmon sperm DNA/protein A-Sepharose beads at 4°C for 3 hr. The precipitated beads were washed with different washing buffers [24]. CHIP-eluted DNA was reversed and extracted with phenol/chloroform. The eluted DNA was analyzed by quantitative Real-Time PCR with the following primers for ChIP [25]:

Primers for the c-FLIP promoter: c-FLIP promoter region #1: forward primer 5’- TTAAATGCCTGCCCCTACTG-3’, reverse primer 5-TAGCACCTCCATCACCACCT- 3’; c-FLIP promoter region #2: forward primer 5’- AAGAGAATCGCTTGAACTAGGAAGG-3’, reverse primer 5’- CTATGGCTTGTGTGACTGAGTATGC-3’; c-FLIP promoter region #3: forward primer 5’-TTACCTTCAGCATCAGGTAGCTAGG-3’, reverse primer 5’- CTCTTGGATCAGAATGTGAGAGTCA-3’

### Western Blot analysis

An equal amount of protein in the cell lysate was loaded in each well and electrophoresed through 4-12% SDS-PAGE then transferred to a PVDF membrane. The primary antibodies indicated in the figures were incubated with the PVDF membrane after transfer in blocking buffer at 4°C overnight. Antibody binding was detected on PVDF with appropriate IR Dye-secondary antibodies (LI-COR Biosciences, USA) by the ODYSSEY infrared imaging system or with ECL Reagent (Thermo Fisher Scientific, catalog no. 32106) chemiluminescence reaction with appropriate horseradish peroxidase (HRP)-conjugated secondary antibodies (Thermo Fisher Scientific, catalog no. no. 31460 for Goat anti-rabbit IgG and catalog no. 31430 for Goat anti-mouse IgG) by the Syngene imaging system.

### c-FLIP promoter-driven luciferase reporter assay

c-FLIP-promoter-luciferase plasmids were generated previously in the lab [26]. The FLIP-promoter-luciferase plasmids were transiently transfected with lipofection 2000 (Life Technologies, catalog no. 11668-027) into cancer cells as stated in the figures for 24 hours, followed with Adenovirus-p73 infection for 24 hours. Luciferase reporter activity in cells was measured based on the bioluminescence using IVIS after addition of luciferin.

### Knockdown of gene expression by siRNA

Cells were transfected with siRNA using lipofectamine RNAiMAX (Life Technologies, catalog no. 13778075) as described in the protocol from the manufacturer. At 48 hr after transfection, cells were further treated as indicated in the figures.

### Flow Cytometry Assay

Briefly, cells were fixed with 70% ethanol, and stained with Propidium Iodide (PI), then subjected for CytoFLEX to measure DNA content of the PI-stained cells. Cell cycle distribution was analyzed with Flowjo.

### Cytokine profiling assay

Cell culture supernatant levels of the cytokine indicated in the supplementary table 1 were measured with a Human Premixed Multi-Analyte Kit (R&D Systems, Inc., Minneapolis, MN) using a Luminex 200 Instrument (LX200-XPON-RUO, Luminex Corporation, Austin, TX) according to the manufacturer’s instructions. Analyte values were reported in picograms per milliliter (pg/mL). Any values that say “<X” mean the sample value for that analyte was below the lower limit of detection, and any value that says “>X” means the sample value for that analyte was above the upper limit of detection. Any values that say “<X” was recoded as zero in this study.

### Statistical analysis

Statistical analyses were performed using the student’s t-test or Anova with GraphPad Prism software. Statistical significance was determined by p<0.05. Combination indices were calculated using the Chou-Talalay method with CompuSyn software.

## Results

### The *c-FLIP* gene is transcriptionally regulated by the tumor suppressor p73 protein

To determine whether p73 regulates c-FLIP-L/S expression in cancer cells, we transiently overexpressed p73-α and p73-β, the two p73 active isoforms, in cancer cells using lipofectamine transfection and adenovirus infection, respectively. Both the overexpression of p73-α and p73-β dramatically increased c-FLIP-L/S at the protein level in various cancer cell lines carrying either wild-type p53 or mutant p53 (**Figure 1A and 1B, Supplementary Figure S1A**). To further examine the c-FLIP-L/S expression in *TP53*-deficient cells as compared to wild-type p53 expressing cells with same genetic background, we generated *p53*-knockout (KO) U2OS using CRISPR and established stable expression of mutant p53-R175H in the U2OS cell line. The knockdown of *p73* expression in these three cell lines reduced c-FLIP-L/S expression at the protein level as compared to siRNA control (**Figure 1C**). The same effect of the knockdown of *p73* on *c-FLIP* expression was also observed in other p53-mutant cancer cells (**Supplementary Figure S1B**). These results, taken together, suggest p73-dependent c-FLIP-L/S expression in cancer cells regardless of p53 status.

**Figure 1.**
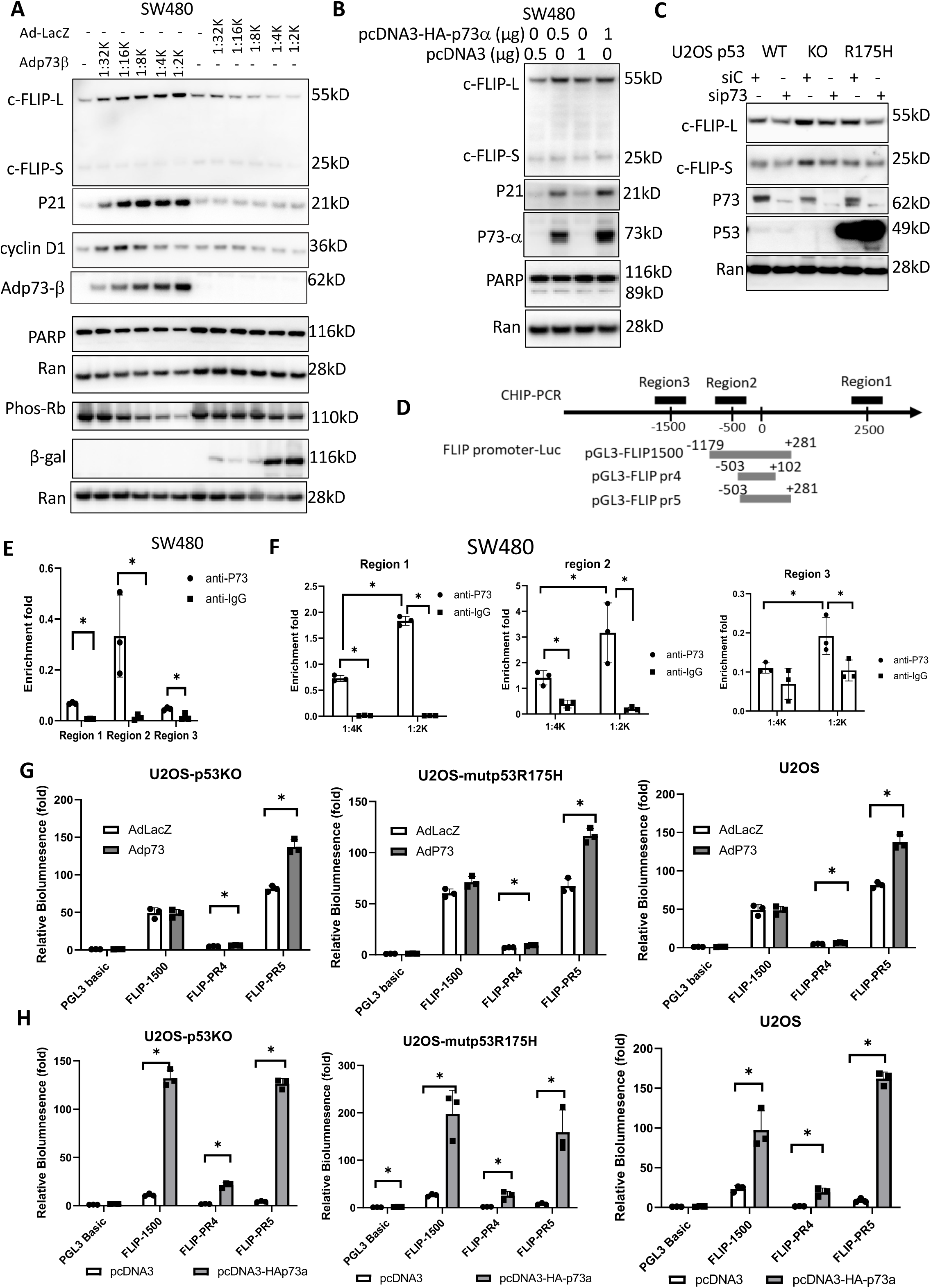
*c-FLIP* is a transcriptional target of p73. A. The overexpression of p73-β increases c-FLIP expression at the protein levels. SW480 cells were infected with recombinant p73-Adenovirus for 48 hr. B. The overexpression of p73-α increase c-FLIP expression. SW480 cells were transfected with pcDNA-HA-p73-α using lipofectamine. C. Knockdown of *p73* reduces c-FLIP protein expression. U2OS cells with p53 wild-type (WT), p53 knockout (KO) or mutant p53 (R175H) were transfected with siRNA targeting *p73*. The c-FLIP protein expression (A, B and C) was examined by Western Blot. D. The fragments flanking the *c-FLIP* promoter used for CHIP-PCR and Luciferase reporter assay. E. Endogenous p73 binding to *c-FLIP* promoter examined by ChIP-PCR assay. F. CHIP-PCR assay for the enrichment *of c-FLIP* promoter bound with Ad-p73 in SW480 cancer cells. The *c-FLIP* promoters were quantified by real-time PCR using the ChIP-eluted DNA. Data were normalized to the input respectively. Data represent mean ± SD. *, P < 0.05 (E and F). G. The *c-FLIP* promoter-luciferase reporter assay in the cells infected with the recombinant adenovirus p73-β for 20 hours. H. *c-FLIP* promoter-luciferase reporter assay in cells transfected with pcDNA-HA-p73-α for 20 hr. Data represent mean ± SD. *, P < 0.05 (G and H). The data (A-C,and G) represents at least two experiments, and the data (E,F and H) represents one experiment. Data are expressed as mean ± SD. *, P < 0.05

To examine if p73 transcriptionally upregulates c-FLIP expression, we performed a ChIP-PCR analysis using the primers flanking three different regions of the *c-FLIP* promoter and further did *c-FLIP* promoter-driven luciferase reporter assays accordingly (**Figure 1D**). The ChIP-PCR assay detected the *c-FLIP* promoter regions bound with endogenous p73 in *p53*-mutant cancer cells (**Figure 1E**) and further revealed an increase in *c-FLIP* promoter bound with the recombinant p73-β in a p73-level dependent manner in the cancer cells (**Figure 1F**). These results, taken together, suggest the binding of p73 to the *c-FLIP* promoter in cancer cells. Further mechanistic studies using *c-FLIP* promoter-driven luciferase reporter assay showed that both p73-β and p73α increased the *c-FLIP*-promoter (PR4 and PR5)-Luciferase activity with the highest PR5- luc upon the p73 overexpression in U2OS, P53-KO-U2OS or mutant-*p53* R175H expressing cancer cells (**Figure 1G and 1H**). PR1500-luc reporter assay showed a significant increase in luciferase activity upon p73α overexpression as compared to the overexpression of p73-β in the cells (**Figure 1G and 1H**). These results, taken together, suggest that p73 transcriptionally upregulates *c-FLIP* expression in cancer cells.

### Reduction of c-FLIP-L/S expression sensitizes cancer cells to p73-induced apoptosis in cancer cells

To assess the p73-dependent cell fate decision between cell survival and death, we performed a cell cycle assay and found that 5-10% of cells were detected in the subG1 phase (**Figure 2A**), consistent with the observation of no increase in cleaved-PARP, a cell death marker, in the cells following p73-β overexpression for 2 days (**Figure 1A**). Moreover, cell cycle analysis showed ∼60% of the p73-overexpressing cells in the G1 phase as compared to the 40% of the control cells (**Figure 2B**). Consistent with the cell cycle profiling, the level of phospho-Rb was reduced, and cyclin D1 was increased, both of which were correlated to increased p21 protein in cells with p73-β overexpression (**Figure 1A**). These results suggest cell survival along with cell cycle arrest in cells upon the p73 overexpression under the experimental conditions. To further examine the effect of p73 on cell death phenotypes, we extended the duration of p73 overexpression from 48 hours to 72 hours in tumor cells and found an increase of cell number in the sub-G1 phase of the cell cycle in cancer cells with the extended duration of p73 overexpression to 72 hours (**Figure 2C**).

**Figure 2.**
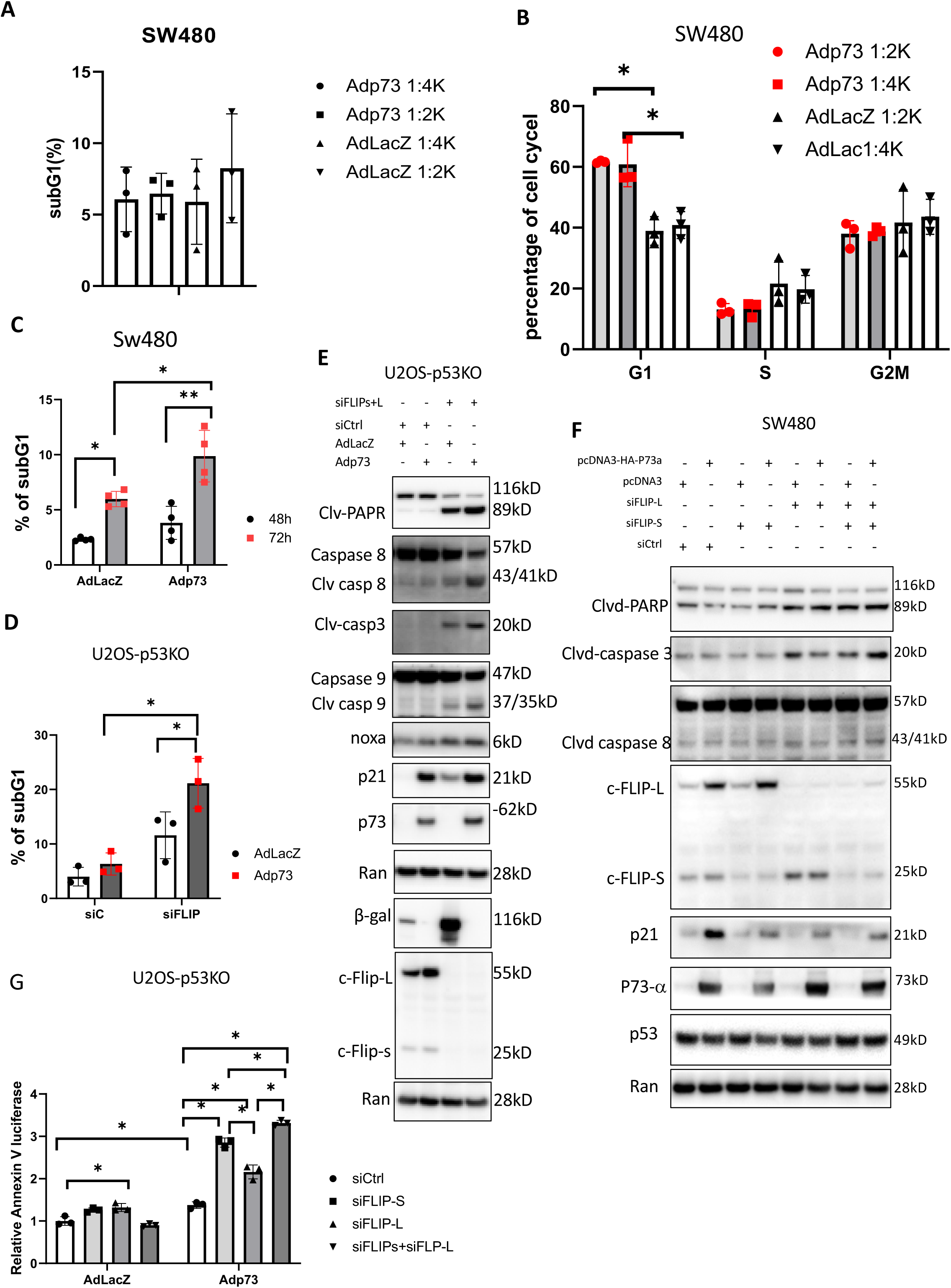
Depletion of *c-FLIP* sensitizes cancer cells to p73-induced cell death. A. Sub-G1 flow cytometric assay in SW480 cells that were infected with Ad-*p73* for 48 hr. B. Cell cycle analysis in SW480 cells infected with Ad-p73 for 48 hr (A) by Flowjo. C. Sub-G1 assay in SW480 cells infected with Adp73 for 48 and 72hours. D. Sub-G1 assay in U2OS-p53KO cells with knockdown of *c-FLIP* and overexpression of *p73-*β*. c-FLIP* was transiently knocked down by siRNA and *p73-*β was overexpressed with adenovirus infection for 30 hr. E. Cleaved-PARP protein in cells with knockdown of *c-FLIP* and overexpression of p73 protein after Ad-*p73* infection for 30 hr. F. Western Blot assay for protein levels in SW480 cells with knockdown of *c-FLIP* isoforms followed by p73α transfection. G. Real time-Glo Annexin V assay. The cells were transfected with siRNA as indicated, followed with AdLacZ or Adp73 infection for 44 hours. The data (A, B and C) represents a combination of two independent experiments. The data (D and E) represents at least two independent experiments. The data (F and G) represent one experiment. Data are expressed as mean ± SD. *, P < 0.05.

To examine if upregulation of c-FLIP inhibits p73-induced cell apoptosis, we knocked down *c-FLIP-L/S* expression in cancer cells. In the knocked-down *c-FLIP-L/S* cells, the overexpression of p73 rapidly increased cell number in the sub-G1 phase at 30 hours (**Figure 2D**), consistent with the enhanced cell death markers cleaved-PARP, cleaved caspase-8 and cleaved caspase-3 in the cells with overexpression of either p73 α or p73 β (**Figure 2E and 2F**). To examine which one of the c-FLIPL/S proteins plays a significant role in attenuating p73-mediated cell apoptosis, we further selectively knocked down *c-FLIP-L* or *c-FLIP-s* using specific siRNA in the cancer cells. The knockdown of *c-FLIP-L or c-FLIP-s* increased PARP cleavage, the cell death marker in cancer cells (**Figure 2F**). Bioluminescent Annexin V analysis showed an increase in bioluminescence in the Adp73-infected cells transfected with siRNAs either targeting c-FLIP-L or c-FLIP-s, or both, indicating cell death (Figure 2G). These results suggest that c-FLIPL/S is a major factor involved in p73-mediated cell fate determination, i.e., c-FLIP- L/S loss promotes p73-induced cell apoptosis.

### p73 induces extrinsic apoptosis in *c-FLIP*-deficient cancer cells

To examine the apoptotic pathway by which p73 induces cell death, we knocked down caspase-8 (a major mediator in the extrinsic pathway) and caspase-9 (a major mediator in the intrinsic pathway) in cancer cells. The knockdown of *caspase-8* or *caspase-9* partially rescued the cell death in the cells with siRNA control in response to the p73 overexpression (**Figure 3A**), suggesting that the longer exposure to p73 activation causes cell death partly via the intrinsic and/or extrinsic apoptotic pathways. By contrast, the effect of the p73 on cell death was blocked mainly by the knockdown of caspase-8 in the *c-FLIP* knockdown cancer cells based on the sub-G1 flow cytometric assay (**Figure 3A**). We further found that knockdown of *caspase-8* blocked PARP cleavage in the *c-FLIP*-knockdown cancer cells in response to either p73-β or p73-α overexpression, but the knockdown of *caspase-9* could not do so in cells with treatment for 30 hours (**Figure 3B, supplementary Figure S2**). These results suggest that p73 induces cell death mainly via extrinsic apoptosis in *c-FLIP*-deficient cells.

**Figure 3.**
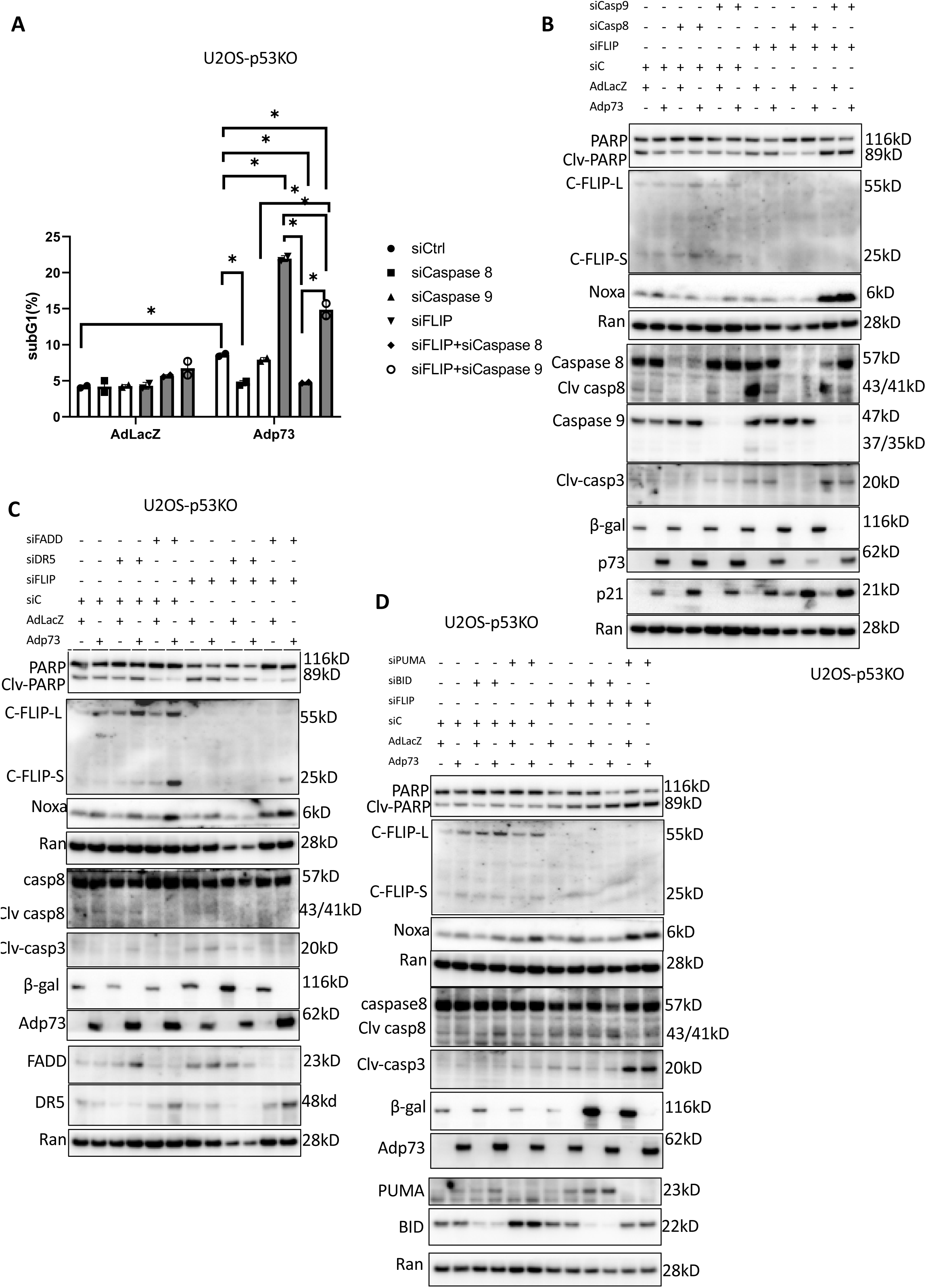
p73 induces cell death via extrinsic apoptosis in *c-FLIP*-deficient cancer cells. A. Sub-G1 analysis in U2OS-p*53*KO cancer cells with knockdown of *caspase-8* and *caspase-9*. The cells were transfected with the siRNAs to knockdown *caspase-8*, *caspase-9* or *c-FLIP-s+L* expression, followed with Ad-*p73* or AdLacZ infection for 72 hr. B. Cleaved PARP in U2OS-P53KO cells with knockdown of *caspase-8* and *9*. *Caspase-8*, *caspase-9* and *c-FLIP-s+L* were knocked down by siRNA in the cells followed with Ad-*p73* or AdLacZ infection for 30 hr. C. Cleaved PARP in U2OS-P53KO cells with knockdown of *DR5* and *FADD*. U2OS-*P53*KO cells were transfected with siRNA to knock down *DR5* and *FADD* expression, followed with Ad-*p73 or AdLacZ* infection for 30 hr. D. Cleaved PARP in tumor cells with knockdown of *PUMA* and *BID*. U2OS-*P53* KO cells were transfected with siRNA to knock down *PUMA* and *BID* expression, followed by Ad-*p73 or AdLacZ* infection for 30 hr. The data (A-D) represents two experiments. Data are expressed as mean ± SD. *, P < 0.05

TRAIL Death Receptor DR5 is an upstream cellular factor in the extrinsic cell death pathway. To further examine if the extrinsic pathway is required in this process with a short expose of p73 overexpression in the *c-FLIP*-knockdown cells, we knocked down *DR5* and *FADD*, the components of the Death-Inducing-Signaling-Complex (DISC). The knockdown of either *DR5* or *FADD* partially blocked PARP-cleavage in the *c-FLIP*- knockdown cancer cells upon p73-β overexpression at 30 hr (**Figure 3C**). By contrast, knockdown of *PUMA*, a factor involved in the intrinsic pathway, could not block p73- induced cleavage of PARP in the *c-FLIP* KD cells (**Figure 3D**). Activation of caspase-8 may result in cleavage of Bid, a Bcl-2 family protein with a BH3 domain only, which, in turn, translocates to mitochondria to release cytochrome c thereby initiating a mitochondrial amplification loop [27]. The pro-apoptotic Bid connects the activation of the extrinsic death receptor pathway to activation of the mitochondrial-disruption processes associated with the intrinsic pathway. The knockdown of *Bid* could not attenuate the cleavage of PARP or caspase-9 in *c-FLIP*-knockdown cells (**Figure 3D**).

To identify if cell death ligands engage in the effect of p73 activation of apoptosis, we further examined cytokine secretion from cancer cells upon p73 overexpression. We transiently transfected *p73-*α into cells using lipofectamine. The cytokine profiling showed an increase of cell death ligands such as TNF and TRAIL in the cell culture media upon the p73-α overexpression in the cancer cells (**Supplementary Table 1**).

### Small-molecule CB-7587351 “switcher compound” alters p73 outcomes by upregulation of proapoptotic signaling and reduction of c-FLIP in p53-mutant cancer cells

We further applied the above insights to search for small-molecule compounds altering outcomes of p73 activation between proapoptotic signaling and c-FLIP levels for cancer therapy. We screened a Chembridge library of small-molecule compounds that might restored p73 signaling via a high-throughput screen and examined c-FLIP protein expression in cancer cells treated with hits from the library. We identified small molecule CB-7587351 (2-[(8-ethoxy-4-methyl-2-quinazolinyl) amino]-5,6,7,8-tetrahydro-4(1H)- quinazolinone) that reduced c-FLIP protein expression in p53-mutant and p53 wild-type cancer cells (**Figure 4A and 4B**). The Real-time PCR assay showed no significant changes of *c-FLIP* expression at the mRNA level in the cancer cells treated with CB- 7587351 (**Figure 4C**), but the western blot assay showed a decrease in c-FLIP-L/S at the protein level in the cancer cells (**Figure 4B and 4D**). Furthermore, CB-7587351 treatment reduced p73-mediated upregulation of c-FLIP protein in cancer cells with overexpression of p73 (**Figure 4D**). To examine if treatment with CB-7587351 reduces c-FLIP protein stability, cancer cells were treated with MG132, a proteasome inhibitor, the treatment with MG132 rescued c-FLIP from the CB-7587351-induced protein degradation in the cancer cells (**Figure 4E**). These results suggest that CB-7587351 induces c-FLIP protein degradation and prompted us to define it as a “switcher compound” as the consequence is an impact on cell fate from cell survival to cell death. We further knocked down *<u>C</u>-terminus of <u>H</u>sc70-Interacting <u>P</u>rotein (CHIP)* and *ITCH*, two E3 ligases that have been reported to be involved in c-FLIP degradation in cancer cells [28, 29]. The knockdown of either *CHIP* or *ITCH* could not rescue c-FLIP from the CB-7587351-induced protein degradation in the cancer cells (**Figure 4F and 4G**). We also treated cells with caspase pan inhibitors or siRNA to silence caspase 8 in cells, and the cells with the inhibition of caspases showed the degradation of c-FLIP in response to the CB7587351 treatment (figure 4F, supplementary figure S3), suggesting that the c-FLIP reduction is not due to cell death at the early time points.

**Figure 4.**
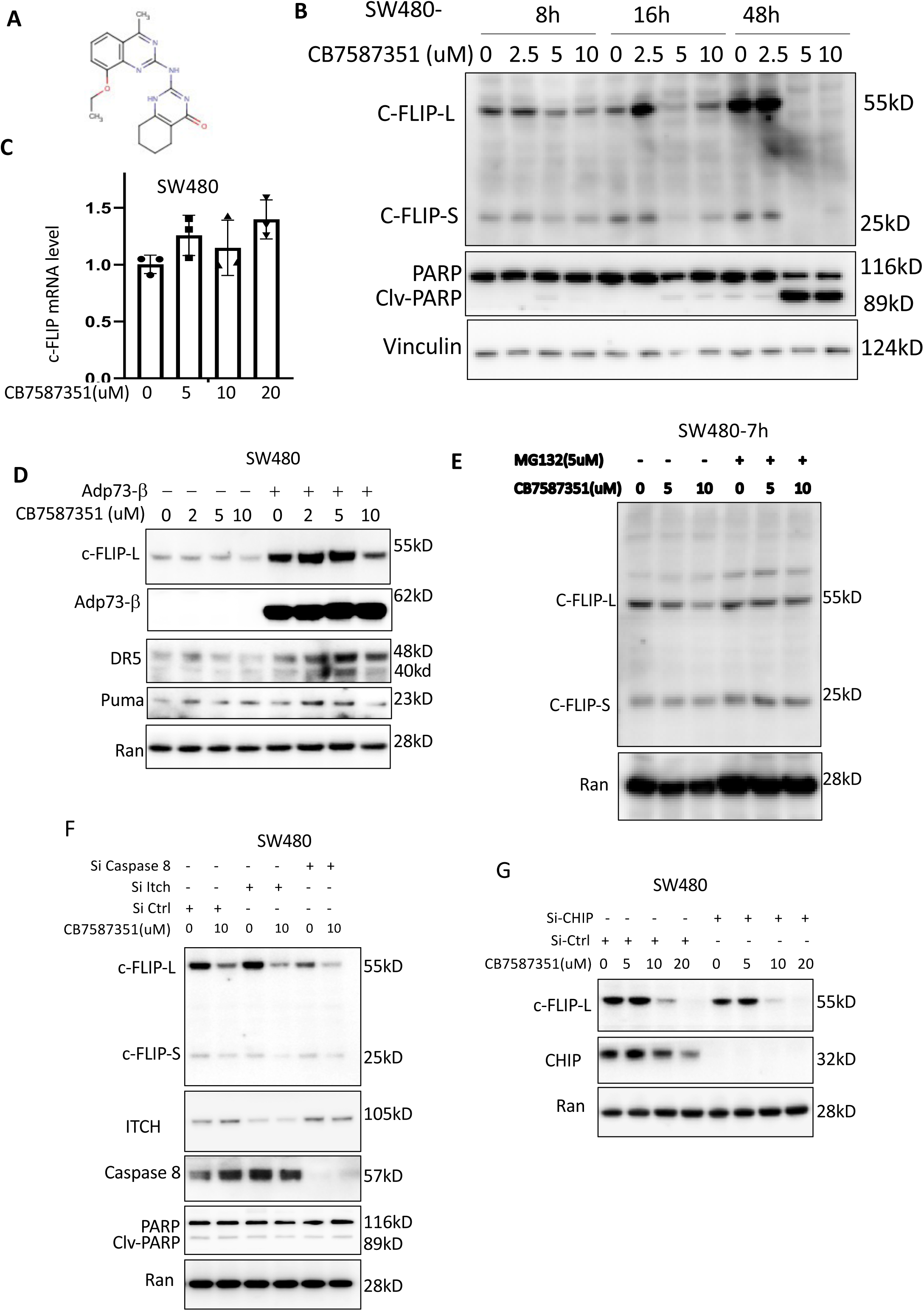
Identification of small molecule “switch compound” CB-7587351 as a c-FLIP degrader. A. Molecular structure of **“switch compound”** CB-7587351. B. c-FLIP protein levels in tumor cells treated with **“switch compound”** CB-7587351. C. mRNA levels of *c-FLIP* in cells treated with CB-7587351. D. Protein levels of c-FLIP in SW480 cells with overexpression of recombinant *p73-*β, followed by CB-7587351 treatment for 8 hr. E. Protein levels of c-FLIP in cells treated with CB-7587351 and MG132. F. c-FLIP protein levels in SW480 cell with knockdown of *ITCH or* caspase 8. The *ITCH* or Caspase 8 was knocked down with siRNA for 48 hr, followed with CB-7587351 treatment for 8 hr. H. c-FLIP protein levels in the *ChIP*-knockdown tumor cells. *ChIP* was knocked down by siRNA in SW480 cells, followed by CB-7587351 treatment for 8 hr. The data (B) represents at least two experiments related to 8- and 16-hour time points. The data (C, D and G) represents one experiment. The data (E and F) represents two experiments about ITCH. Data are expressed as mean ± SD. *, P < 0.05.

In addition to the above observations, small molecule CB-7587351 restores p53 pathway signaling in cancer cells as a *bone fide* p53 pathway restoring compound (**Figure 5**). CB-7587351 strongly induced p53-responsive bioluminescence in different cell lines SW480 (mutant *p53* R271H and P309S), DLD1(mutant p53 S241F) and HCT116 p53-/-colorectal cancer cells as well as in wild type *p53*-expressing cancer cells (HCT116) in a dose dependent manner regardless of p53 (**Figure 5A**). Endogenous p53 targets were increased at the protein level by CB-7587351 treatment in *p53*-mutant and wild colorectal cancer cell lines (**Figure 4D**, **Figure 5B**). A real-time RT-PCR assay showed a slight increase of the selected p53 targets at the mRNA level in the cells treated with CB-7587351 (**Figure 5C**). These results suggest that CB-7587351 restores p53 pathway signaling in cancer cells carrying wild-type *p53* or mutant *p53*. To examine if *p73* is involved in the CB-7587351-reactived *p53* pathway in mutant *p53*-expressing cancer cells, we overexpressed recombinant *p73-*β with adenovirus infection and observed the effect of CB-7587351 on the PG13-luc reporter to be further enhanced by the overexpression of p73-β as well as wild-type p53 (**Figure 5D and 5E**). The knockdown of *p73* blocked the activation effect of CB-7587351 on p21 and puma expression in the DLD-1 cells (**Figure 5F**). The transient transfection of siRNA-targeting p73 attenuated the CB7587351-induced PG-13-lucifease reporter in p53-KO cancer cells (**Figure 5G**). These results suggest that CB-7587351 restores p53 pathway signaling via activation of p73 in *p53*-mutant or *p53*-null cancer cells.

**Figure 5.**
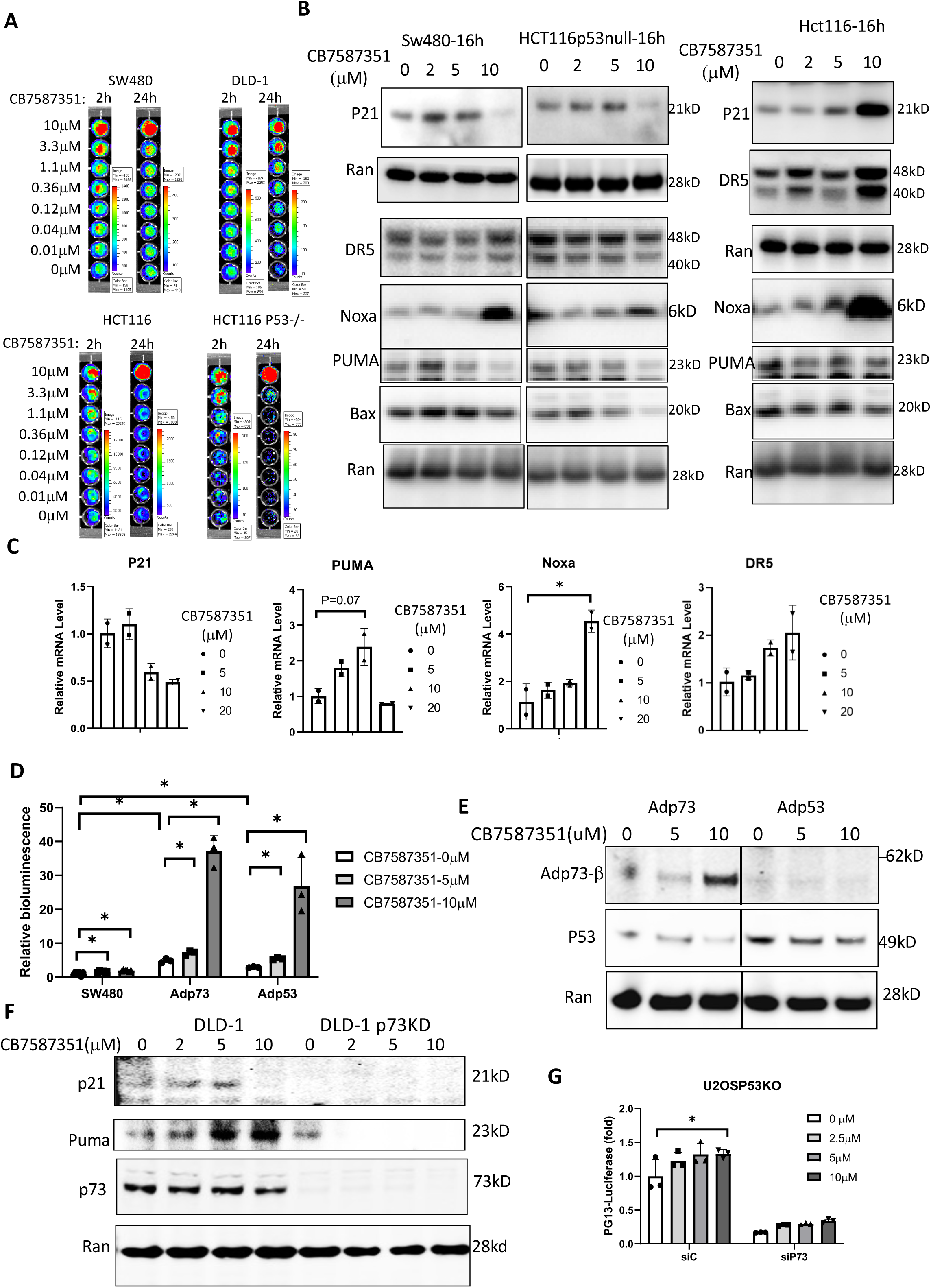
CB-7587351 activates p73-dependent pro-apoptotic transcriptional target genes. A. PG13-bioluminescent assay in cancer cells treated with CB7587351 at different time points. B. The protein levels of p53 targets in different cancer cells treated with CB-7587351 for 16 hr. C. mRNA levels of p53 target genes in SW480 cells treated with CB-7587351 for 3 hr. mRNAs were examined by real time PCR. Data was normalized to GAPDH expression and plotted relative to cells treated with DMSO as control. Data are expressed as mean ± SD. *, P < 0.05 vs. control. D. PG13-luciferase reporter assay in cells with overexpression of *p53* or *p73*. Cells were infected with Ad- *p53* or Ad-*p73*, followed by CB-7587351 treatment for 16 hr. Relative bioluminescence was normalized to those of DMSO treatment in the cells without the infection. Data are expressed as mean ± SD. * P < 0.05. E. The expression of Adp53 and Adp73 in the cells (D). All lanes are from the same western blot. The lane 3 and lane 4 were spliced together from the same blot. A vertical black line indicates the splicing position. The lane (20uM) between Lane 3 and lane 4 was removed because this high dose caused potential side effects at this time point, and it is not suitable for the luciferase analysis. F. Protein levels of p53 targets in *p73* knockdown DLD-1 cells treated with CB-7587351 for 8 hr. G. PG13-Luciferase reporter assay. Cancer cells were transfected with siRNA targeting p73, followed with CB7587351 (μM) treatment. The relative Bioluminescence levels were normalized to those of siCtrl + DMSO. The data (A, C, D, E and F) represent one experiment. The data (B) represents at least two independent experiments. The data (G) represents four independent experiments. Data are expressed as mean ± SD. *, P < 0.05.

We examined whether DNA damage engages in the mechanism of the action of CB- 7587351 on p53 signaling restoration (**Supplementary Figure S4**). The foci and levels of the phosphorylation of γ−H2AX, a sensitive indicator for DNA damage due to DNA double-strand breaks, were measured in the cancer cells. No phosphorylation of γ- H2AX was found in SW480 cells in response to an increased dose of CB-7587351 during early time points following CB-7587351 treatment (**Supplementary Figure S4A, S4B and S4**C). Moreover, no DNA-intercalation was detected (**Supplementary Figure S4D**). These results, taken together, suggest little or no genotoxicity of CB-7587351.

### CB-7587351 induces extrinsic apoptosis in mutant-*p53* expressing cancer cells

To evaluate the therapeutic index of CB-7587351, we examined cell viability of a panel of cancer cell lines with CB-7587351 treatment. The IC50s of CB-7587351 were much lower in cancer cells than those in normal fibroblast MRC-5 cells (**Figure 6A**), indicating a favorable therapeutic index for CB-7587351. A flow cytometric assay showed an increase in sub-G1 content in cancer cells, but not in normal fibroblast cells MRC-5 after CB-7587351 treatment (**Figure 6B**). In addition, CB-7587351 treatment dramatically reduced colony formation in colorectal cancer cells (**Figure 6C**). Knockdown of *p73* abrogated the effect of CB-7587351 on sub-G1 content and PARP cleavage in DLD-1 cells (**Figure 6D and 6E**).

**Figure 6.**
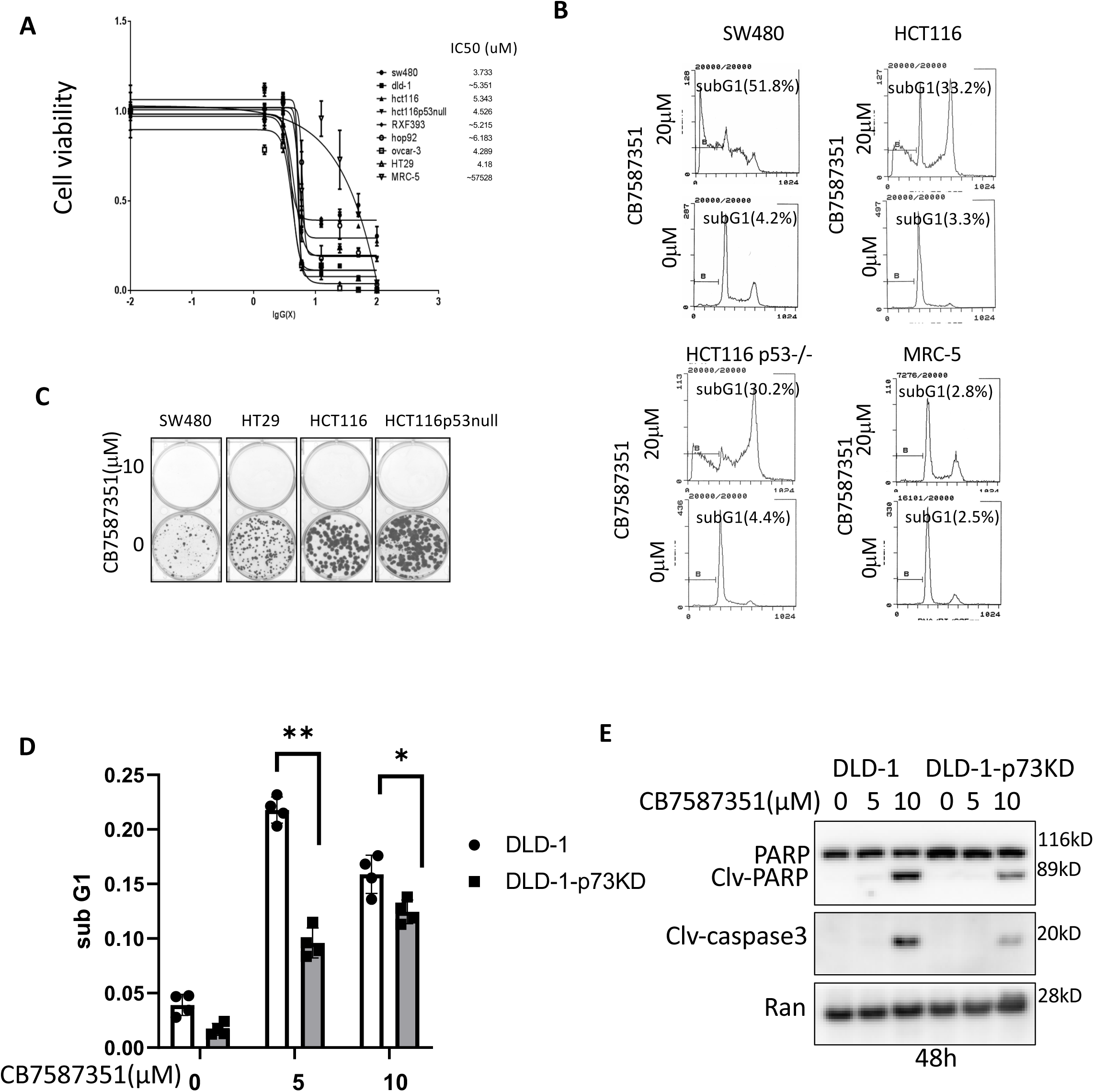
Pro-apoptotic effects of CB-7587351 *on cancer cells.* A. Cell viability assay. Cells were treated with CB-7587351 for 48 hr. B. Cell cycle profiles in the cells treated with CB07587351 for 48 hr. C. Colony formation assay as indicated. D. Sub-G1 assay in *p73*-knockdown DLD-1 cells treated with CB-7587351 for 72 hr. E. Protein analysis of cleaved PARP and cleaved caspase 3 in *p73*-knockdown DLD-1 cells treated with CB-7487351 for 48h. The data (A, B, C) represent one experiment. The data (D and E) represent two experiments. Data are expressed as mean ± SD. *, P < 0.05.

An increase of cleaved caspase-3 and caspase-3/7 activity was detected in cancer cells treated with CB-7587351 at early time points (**Figure 7A and 7B**). Treatment with z-VAD-FMK, a pan-caspase inhibitor, blocked the effect of CB-7587351 on apoptosis in SW480, HT29, HCT116 *p53*-/- and HCT116 colorectal cancer cells according to the sub-G1 assay (**Figure7C and 7D**). We further examined which one, the extrinsic or intrinsic apoptotic pathway, is involved in the induction of the apoptosis. Knockdown of *caspase-8* prevented apoptosis induced by CB-7587351 at 10 μM based on the lack of cleavage of PARP and Annexing-V levels, but knockdown of *caspase-9* could not (**Figure 7E and 7F).** These results suggest that CB-7587351 induces extrinsic apoptosis in *p53*-mutant cancer cells. Knockdown of caspase 9 showed a lesser extent of cleaved caspase 3 as compared to the siRNA control in the cells treated with CB7587351(**Figure 7E**). These results suggest that other caspases might also be involved in the process.

**Figure 7.**
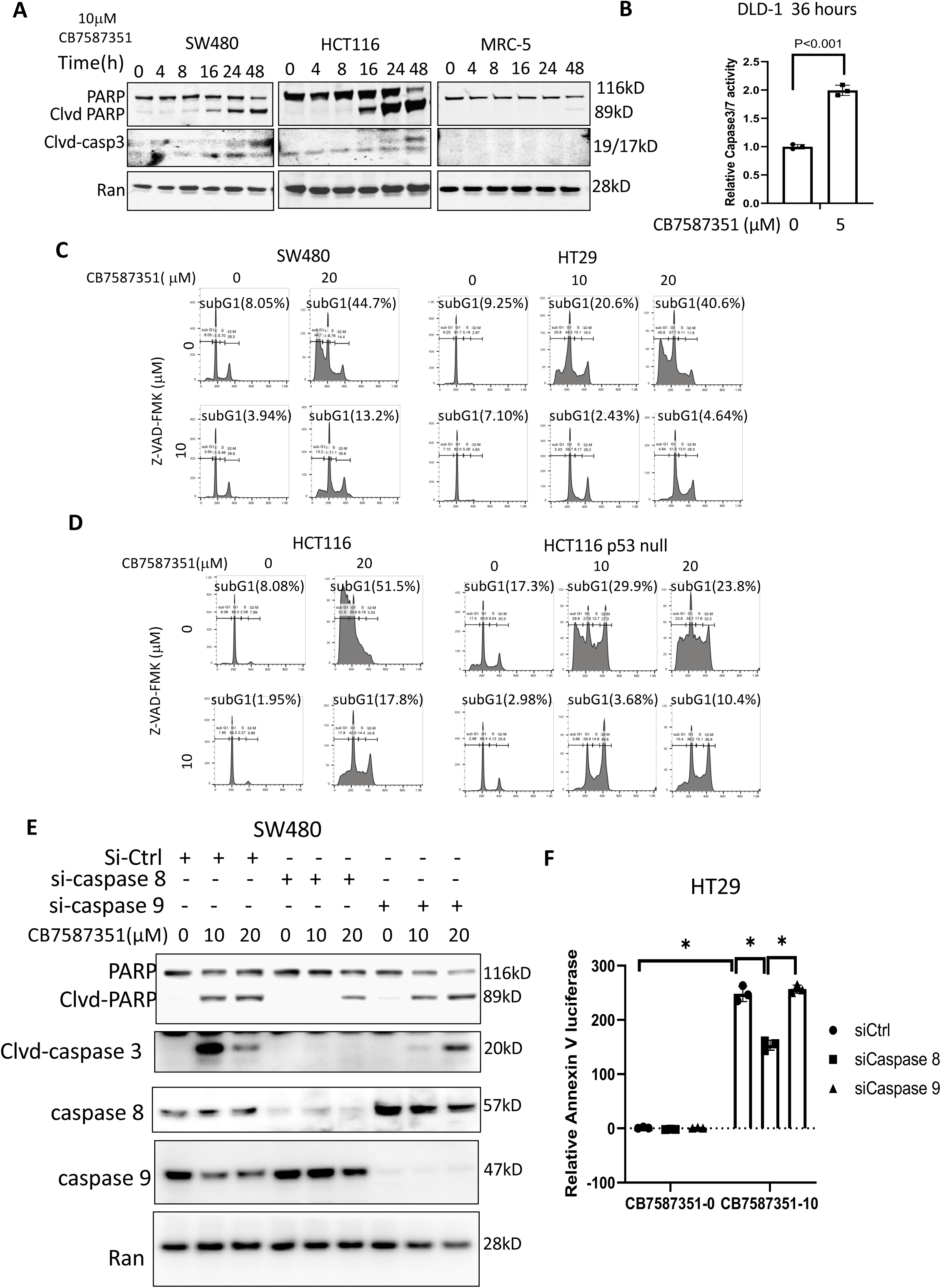
CB-7587351 induces cell death via the extrinsic apoptotic pathway. A. Cleaved caspase-3 in the cells treated with CB-7587351. B. Caspase-3/7 activity in DLD-1 cancer cells treated with CB-7587351 for 36 hours. C. Cell cycle profiles in *p53*- mutant cancer cells treated with CB-7587351 and pan-caspase inhibitor z-VAD-FMK. D. Cell cycle profiles in *p53* wild-type and *p53*-null cancer cells treated with CB-7587351 and pan-caspase inhibitor z-VAD-FMK. E. Cleaved PARP in cancer cells with knockdown of *caspase-8* or *caspase-9*, followed by CB-7587351 treatment. F. Bioluminescent Annexin-V assay in HT29 cells. The cells transfected with siRNA to knock down caspase 8 or caspase 9, followed with CB-7587351(μM) treatment for 41 hours. The data (A, C, D and F) represents one experiment. The data (B and E) represents two independent experiments. Data are expressed as mean ± SD. *, P < 0.05.

As TRAIL is a major death ligand driving cell death via extrinsic apoptosis and the innate immune system, we observed that CB-7587351 treatment synergized with TRAIL to reduce cell viability in cancer cells, but not in normal cells at the tested doses (**Figure 8A and 8B**). CB7587351 treatment enhanced TRAIL-induced PARP cleavage in cancer cells (**Figure 8C, Supplementary Figure S5**). The combinational effect on PARP cleavage was blocked by knockdown of *caspase-8* (**Figure 8D**). These results suggest that CB-7587351 sensitizes cancer cells to TRAIL-induced cell apoptosis via extrinsic apoptosis.

**Figure 8.**
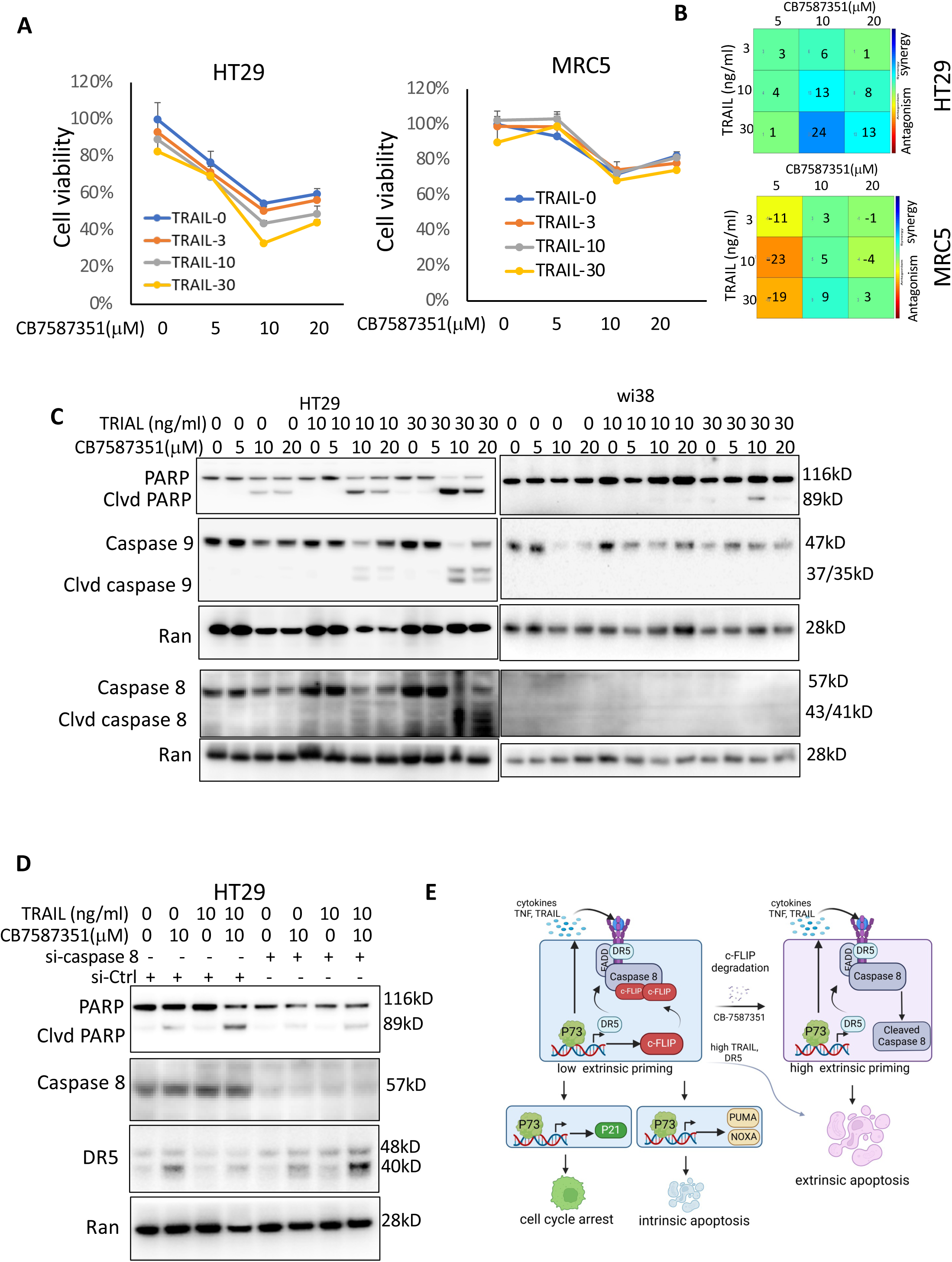
Combinational treatment of TRAIL and CB-7587351 in cancer cells reveals synergy. A. Cell viability assay. B. The synergy assay (A). C. Protein assay of cleaved PARP in tumor cells treated with CB-7587351 and TRAIL. D. Cleaved PARP in HT29 cells with knockdown of *caspase-8*, followed by the combinational treatment. E. The schematic of the mechanism of p73 priming of extrinsic apoptosis in cancer cells. Created with Biorender.com. The data (A, B and D) represents one experiment, and the data (C) represents at least two independent experiments.

On the basis of these findings, we propose that CB-7587351 “switch compound” alters the p73 outcome cell fate balance via degradation of c-FLIP, promoting cells to undergo p73-primed extrinsic apoptosis (**Figure 8E**).

## Discussion

Current approaches for targeting p73 are mainly focused on activating p73 proapoptotic signaling, which can compensate for wild-type p53 deficiency in *p53*-mutant cancer cells, to induce cell death for cancer therapy [9–12]. However, outcomes of p73 activation are complex, and this can influence the anti-tumor efficacy of p73-targeting therapeutic approaches. There is a clear need to understand mechanisms by which p73 determines cell fate between growth arrest (cell cycle arrest or senescence), pro-survival phenotypes, or cell death including apoptosis. We demonstrate here that p73 transcriptionally activates the *c-FLIP-L/S* gene that encodes an an anti-apoptotic factor. The upregulation of *c-FLIP-L/S*, in turn, results in a low level of p73-priming in extrinsic apoptosis that promotes cell survival rather than cell death. Targeting c-FLIP-L/S protein degradation with the “switcher compound” approach we describe leads cancer cells towards a p73-induced extrinsic apoptotic cell fate.

p73-mediated transcriptional upregulation of *c-FLIP-L/S* promotes cellular tolerance to p73-activated apoptotic signaling, given that the knockdown of *c-FLIP-L/S* sensitizes cells to p73-induced apoptosis (**Figures 2 and 3**). Like p73, p53 activation has been reported to upregulate *c-FLIP-L*, which prevents p53-induced apoptosis [22]. Paradoxically, while p73 upregulates c-FLIP protein expression, p73 also increases DR5 expression and TRAIL cell death ligand secretion, key factors activating the extrinsic apoptotic (**Figure 3**). Therefore, we hypothesize that p73 primes the extrinsic apoptotic pathway via a cell death ligand-receptor axis, and the priming level is titrated by the upregulation of c-FLIP-L/S (**Figure 8E**).

The extrinsic apoptotic pathway involves engagement of multiple death receptors through formation of a DISC which contains core cell death signaling components such as FADD, DR5 and caspase-8. The DISC initiates a cascade activating caspase-8 and downstream caspase-3 when the receptors are bound with different death ligands such as tumor necrosis factor (TNF) and Fas [30]. Both DR5 and FADD are required for p73 to induce cell death in our experiments with *c-FLIP*-knockdown cells generated during our study (**Figure 3**), indicating that p73 primes extrinsic apoptosis but attenuates it through c-FLIP. Moreover, the cell death ligands, TNF and TNF-related apoptosis-inducing ligand (TRAIL) are secreted by cells following overexpression of p73 (**Figure 3E**), in accordance with the previous reports that have shown that p73 activation increases expression of TNF family members in cells [31]. Like p73, p53 has been reported to increase Fas expression and enhance levels of Fas at the cell surface [32]. In addition, p21(WAF1), the target of p73 and p53, can also increase expression of cytokines at mRNA levels including Fas [33]. These discoveries suggest a potential autocrine regulation of ligands through CD95 (APO-1/Fas) or TRAIL receptor (DR5) involved in p73-priming of extrinsic apoptosis. Death ligands such as TNF or TRAIL promote activation of the initiator caspase-8 through DISC formation that can propagate the apoptotic signal [34]. As c-FLIP-L/S has sequence homology to caspase-8 and caspase-10 but lacks catalytic activity, c-FLIP-S recruitment to the DISC blocks caspase-8 or caspase-10 caspase activation [35]. c-FLIP-L is a pseudo-caspase and regulates caspase-8 activity based on their ratio in forming heterodimers at the DISC [36]. High levels of c-FLIP-L lead to predominant heterodimer formation with caspase- 8 and inhibits caspase-8 activation and apoptosis [19, 36]. Therefore, p73-transcrtiptional upregulation of *c-FLIP-L/S* may impede p73-primed extrinsic apoptosis via inhibition of caspase-8 activation (**Figure 8E**). Indeed, knockdown of *c-FLIP* increases caspase 8 cleavage in response to p73 overexpression (**Figure 2 and 3**). The upregulation of c-FLIP-L and c-FLIP-S attenuates p73 priming of extrinsic apoptosis, (**Figure 2F and 2G**). The poor p73-priming of extrinsic apoptosis (attenuated by c-FLIP) along with cell cycle arrest (regulated by p21(WAF1) that is induced by p73) can confer cells to endure harsh conditions of stress and survive in response to p73 activation. Indeed, the upregulation of *c-FLIP* correlates with p21 expression in response to p73 activation (**Figure 2**), and depletion of *c-FLIP-L/S* increases extrinsic apoptosis correlated with reduced expression of p21 in cells with p73 overexpression (**Figure 2**). These results suggest that the high priming level of extrinsic apoptosis converts to cell death rather than survival (cell cycle arrest) upon p73 overexpression (**Figure 8E**) in a *c-FLIP-L/S*-dependent manner.

Intrinsic apoptosis appears to be another predominant mechanism by which p73 induces cell death. The signaling involved in intrinsic apoptosis is enhanced by p73 overexpression in tumor cells (**Figure 3D**), but no or less cell death was detected at early time points of 30-48 hr (**Figure 2**). Intrinsic apoptosis occurs through mitochondrial, and mitochondrial apoptosis priming is another key determinant of cell fate [37]. Extended duration (72 hr) of p73 overexpression increased cell death partially via the intrinsic apoptotic pathway in cancer cells with the poor levels of p73-primed extrinsic apoptosis (as compared to the shorter exposures of 30-40 hr) (**Figure 3A and supplementary figure S2**). By contrast, p73 rapidly induces cell death via extrinsic apoptosis in cancer cells when the priming level of extrinsic apoptosis is increased by knockdown of *c-FLIP* (**Figure 2D-2G**, **Figure 3**). We did not measure mitochondrial apoptosis priming in this study. We hypothesize that the priming levels of both intrinsic and extrinsic pathways may have thresholds, and that the cellular apoptotic pathway used is determined by which threshold is breached through first. Depletion of *c-FLIP* may lower the threshold, resulting in highly primed cells readily undergoing p73-induced extrinsic apoptosis. The poor priming of extrinsic apoptosis by the upregulation of *c-FLIP* can protect cells from p73-primed and autocrine cell death ligand-driven extrinsic apoptosis. p73 signaling and cell fate can be manipulated with enhanced intensity and extended duration to overcome the c-FLIP-mediated cell survival threshold.

We demonstrate that p73 binds to the promoter of *c-FLIP* and transcriptionally upregulates *c-FLIP* expression in cancer cells (**Figure 1**), thus raising the interesting and important question of how p73 transcriptionally activates c-FLIP. There are multiple transcription factors involved in regulating *c-FLIP* expression, performing as either activators (such as NFκB and p53) or suppressors (such as c-Myc) [22]. The promoter regions tested in this study have been reported by us previously to be bound by c-Myc [26]. Interestingly, p73 has been found to directly interact with c-Myc to regulate gene expression [38, 39]. Whether p73 interrupts c-Myc’s suppressive effect on c-FLIP expression should be examined in the future. Other potential transcriptional cofactors might also impact the function of p73 in regulating *c-FLIP* transcription in these regions, because the p73-β increases PR4- and PR5-luc reporter activities much more than what was observed with FLIP-1500-luc reporter (**Figure 1**). The detailed regulatory mechanisms by which p73 transcriptionally upregulates *c-FLIP* expression will be need to be more thoroughly addressed in the future, and will be helpful for pharmacological intervention targeting c-FLIP related transcription factors accordingly to increase p73- primed extrinsic apoptosis.

Our study indicates that *c-FLIP* is a critical factor for cell fate decisions in p73-primed extrinsic apoptosis. The depletion of *c-FLIP* using siRNA and the small-molecular “switch compound” CB-7587351 results in a high level of p73-primed extrinsic apoptosis, which sensitizes cancer cells to either p73 activation or TRAIL treatment (**Figure 2 and 8**). Because of the homology between *caspase-8* and *c-FLIP*, inhibiting c-FLIP activity is a big challenge. Targeting c-FLIP degradation via E3 ligase-mediated proteolytic ubiquitination appears to be a promising approach for cancer therapy. CB-7587351 identified in this study provides an example of a small molecule with a dual function by depleting c-FLIP and activating p73 signaling to increase anti-tumor efficacy in *p53*- mutant cancer cells, though the specific immediate molecular targets of CB-758735 remain unclear. Given the effects of CB-7587351 on both p53/p73 activation and c-FLIP degradation, we speculate that the mechanism of action of CB-7587351 on the activation of p73 may, in part, share some signaling pathways involved in degradation of c-FLIP. The detailed molecular mechanism should be investigated more in the future.

c-FLIP is well-known to have high expression in various types of cancer, which is correlated with poor clinical outcomes. In our study, we report dynamic changes of *c-FLIP* expression as a transcription target gene of p73, which results in cell survival following p73-targeting therapy. The role of c-FLIP in p73-mediated determination of cell fate can be exploited more for therapeutic targeting of p73 in cancer therapy. Our study provides an attractive novel approach for targeting c-FLIP-L degradation by “switch compound” CB-7587351 to achieve anti-tumor efficacy through p73-targeting cancer therapy and exploitation of cell fate decisions downstream of p73 signaling leading to apoptosis.

## Acknowledgements

W.S. El-Deiry is an American Cancer Society Research Professor. The work was supported, in part, by the ACS and NIH grant CA176289 to W.S. El-Deiry.

---

**Supplementary Figure S1.**
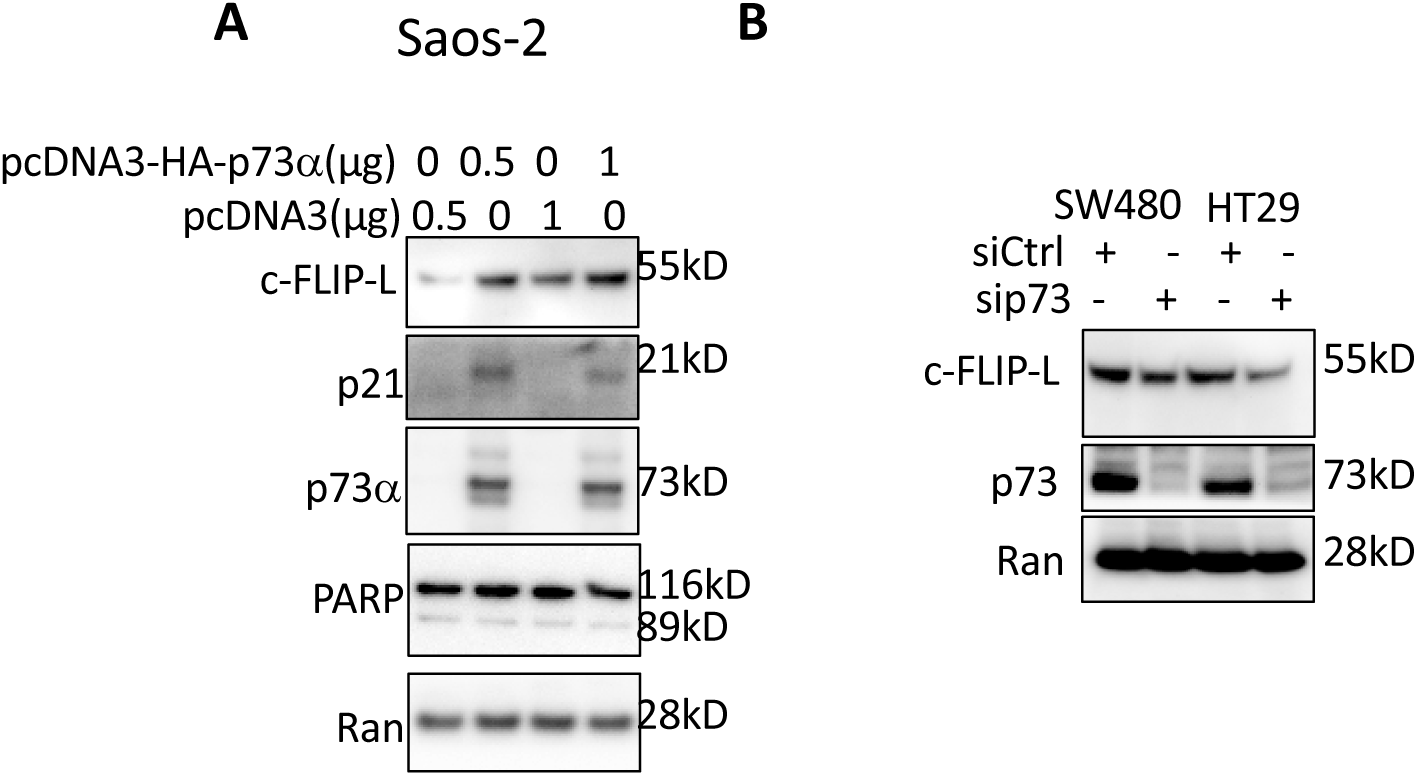
p73 increases *c-FLIP* expression. A. Saos2 cells were transfected with pcDNA3-HA-*P73*α by lipofectamine. B. *p73* expression was knocked down by siRNA in *p53*-mutant cancer cells.

**Supplementary Figure S2.**
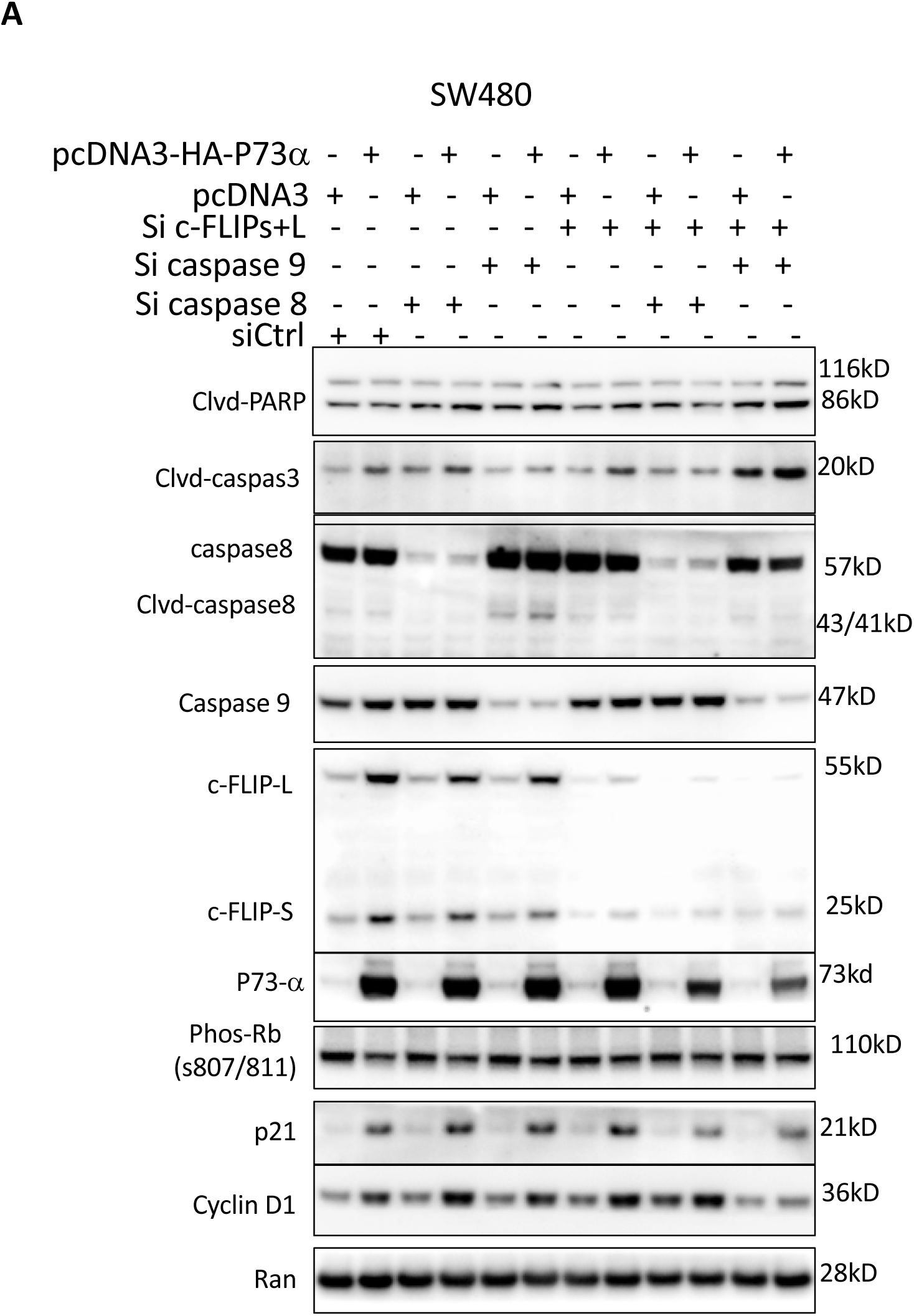
p73α induces extrinsic apoptosis in *c-FLIP*-knockdown cancer cells. A. Knockdown of *caspase-8*, *caspase-9* and *c-FLIP* in SW480 cells, followed with pcDNA3-HA-*p73*α transfection.

**Supplementary Figure S3.**
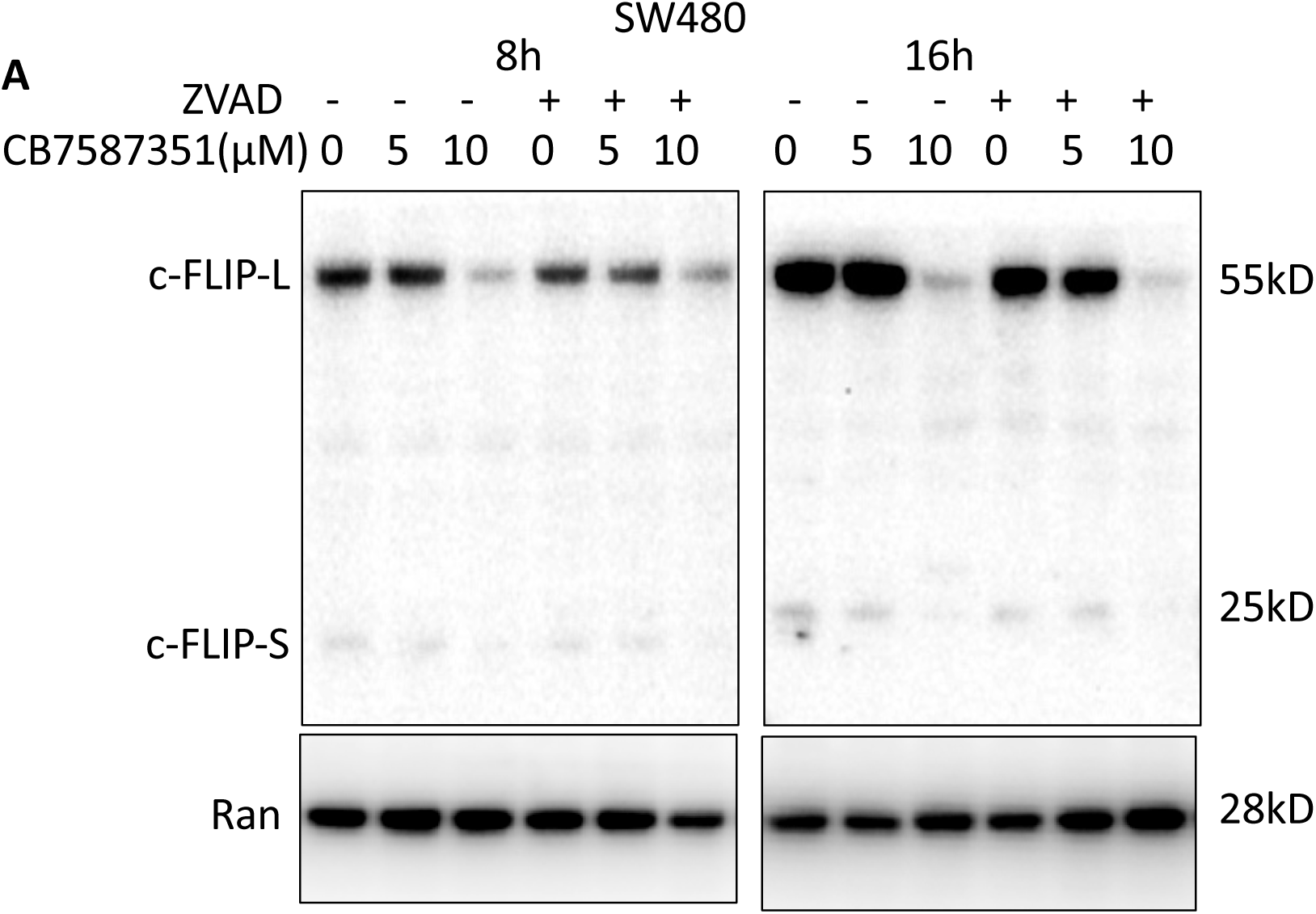
Small molecule CB-7587351 reduces c-FLIP at protein levels. A. SW480 cells were treated with CB-7587351 and pan-caspase inhibitors.

**Supplementary Figure S4.**
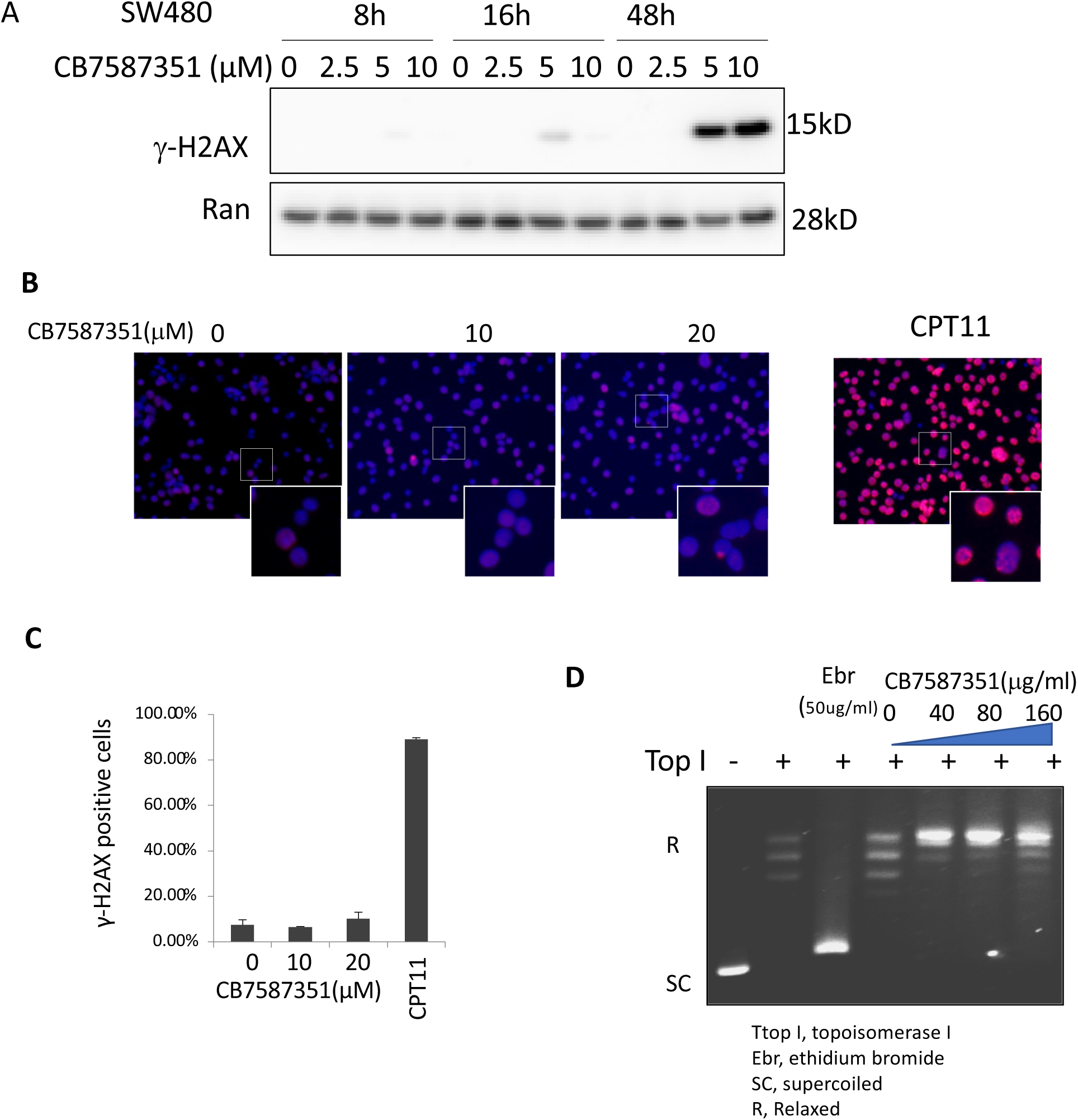
Genotoxicity assay of CB-7587351 in cancer cells. A. Genotoxicity of CB-7587351 was measured by Phosphorylation of H2AX. SW480 cancer cells treated with CB-7587351 at different concentrations over a time course. B. γ-H2AX foci in cancer cells treated with CB-7587351, or CPT-11 as a positive control for H2AX foci staining. C. Number of DNA-damaged cells (B). D. DNA intercalation assay.

**Supplementary Figure S5.**
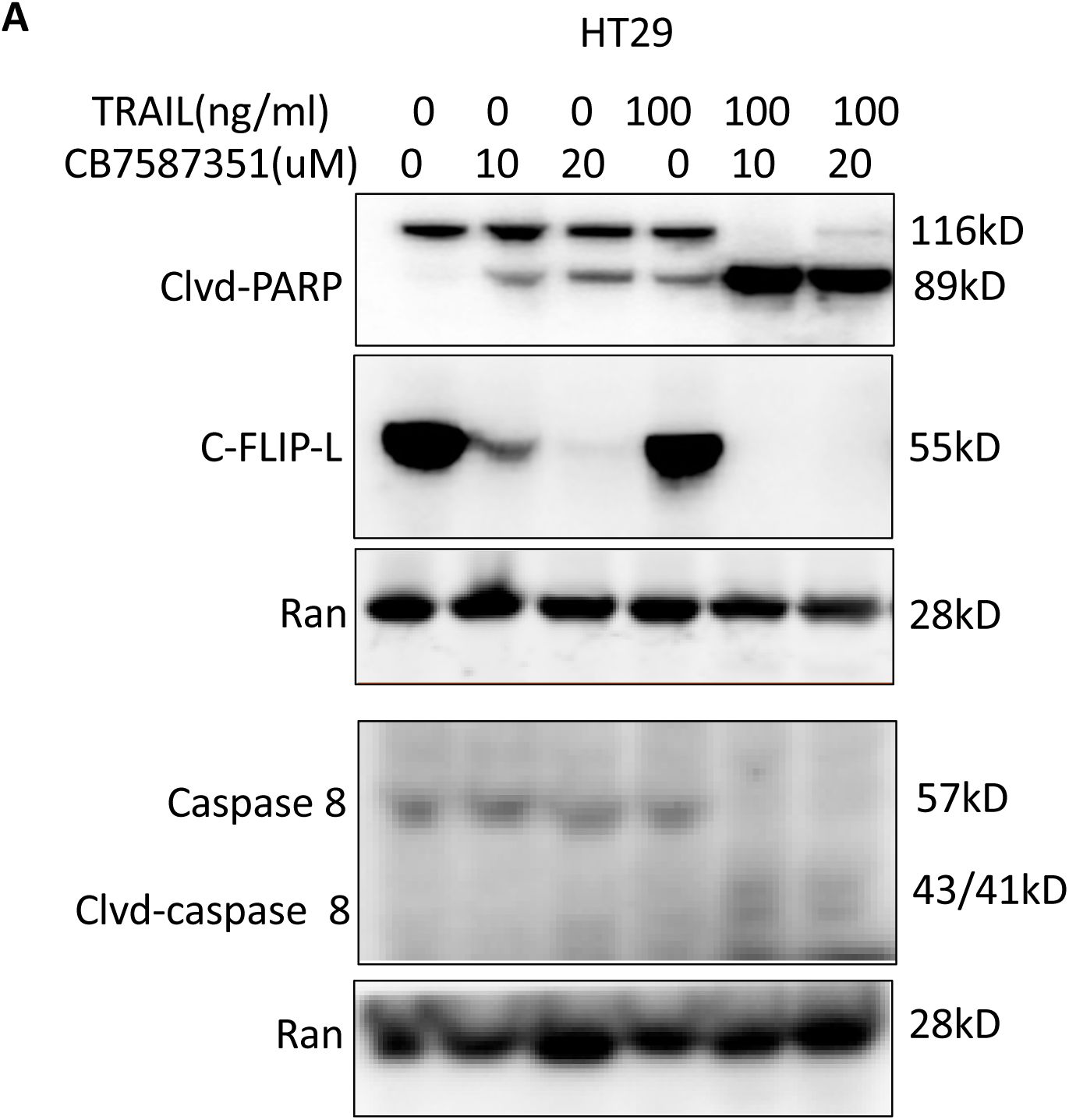
Combinational treatment of CB-7587351 and TRAIL in HT29 cells. A.HT29 cells were treated with CB-7587351, followed by TRAIL treatment. Protein levels were measured by Western Blot.

